# Robust Cable Models from Complex Neuronal Morphologies

**DOI:** 10.64898/2026.05.10.721501

**Authors:** Jordan M. R. Fox, Brian J. Fischer, William DeBello, Jose L. Peña

**Affiliations:** Dominick P. Purpura Department of Neuroscience, Albert Einstein College of Medicine; Zuckerman Institute, Columbia University; Department of Mathematics, Seattle University; Center for Neuroscience, University of California, Davis

**Keywords:** neuron cable model, neuronal morphology, SWC, skeletonization, toric spines, mean curvature flow, multicompartmental simulation, surface mesh

## Abstract

We present a free and open-source, semi-automated, topologically robust pipeline for fitting cable models to 3D surface mesh morphology data of neuronal membranes, particularly suited to structures with complex shapes and topological holes. The motivation for this work is the discovery of morphologically complex neuronal spines on the auditory space-specific neurons of the barn owl, dubbed “toric spines”, notable for their high curvature, branching density, and topological holes. Multicompartmental simulation software requires morphology to be represented as cable models, yet existing software tools for fitting cable models to complex 3D surface meshes generally assume acyclic topology, and thus did not work well for toric spines. We present the Mesh and Skeleton Cable Fitting (MASCAF) pipeline and software, which fits a cable model to a closed 3D surface mesh using mean-curvature flow skeletonization, and automatically handles cyclic topology. In this paper, we demonstrate how MASCAF is applied to fit cable models, how loop connections can be reconstructed in multicompartmental simulation software, and how the results can be validated using geometry and simulator-based methods. While non-tree morphologies such as toric spines are neuroanatomically special, our software pipeline provides a cable-model fitting approach for surface mesh data that is topologically robust, deterministic, open-source, and applicable to general morphologies, thereby closing a crucial gap between neuronal imaging and high-resolution simulation.

## 1 Introduction

### 1.1 From Imaging Data to Multicompartmental Simulation

Neuronal morphology plays a fundamental role in shaping electrical signaling, synaptic integration, and neural computation (Rall 1962; Mainen and Sejnowski 1996; London and Häusser 2005). Since the foundational work of Rall and others on cable theory (Rall 1959), it has been well established that dendritic geometry strongly influences input resistance, voltage attenuation, temporal filtering, and nonlinear integration (Agmon-Snir et al. 1998; Stuart et al. 2016; Koch 1999). Moreover, advances in imaging technologies, particularly serial block-face scanning electron microscopy (SBEM) (Denk and Horstmann 2004) and stimulated emission depletion (STED) microscopy (Hell and Wichmann 1994), now allow reconstruction of neuronal structures at nanometer-scale resolution.

While imaging data and mesh representations preserve detailed geometric features, they are not directly compatible with modern multicompartmental simulators such as `NEURON` (Carnevale and Hines 2006), `GENESIS` (Bower and Beeman 1998), `Arbor` (Abi Akar et al. 2019), `MOOSE` (Dudley et al. 2022), `Brian 2` (Stimberg et al. 2019), `EDEN` (Panagiotou et al. 2022), and `Jaxley` (Deistler et al. 2025), which can provide valuable insights into the relationship between neuronal morphology and function. These simulators, using the well-established cable equations (Rall 1959; Carnevale and Hines 2006), represent neurons as graphs of isopotential compartments, with morphology typically specified by a cable model composed of connected cylindrical or frustum-shaped segments. In such frameworks, realistic morphological specification is important for accurate simulation results.

Fitting cable models, either directly to imaging data or to surface mesh data derived from imaging, is a general challenge (Wang et al. 2020; Liu et al. 2016; Peng et al. 2014). A variety of software tools have been developed for these tasks (Oxford Instruments; Peng et al. 2014; Sato et al. 2000; Acciai et al. 2016), but naturally, some approaches have limitations that make them inappropriate for particularly complex topologies. In this work, we present a semi-automated, topologically robust pipeline for fitting cable models to 3D surface mesh data, without restricting them to tree-like structures.

The central calculation performed by multi-compartmental simulators like `NEURON` and `Arbor` is the construction and integration of the cable equations, a system of first-order nonlinear ordinary differential equations that describe changes in membrane potential for the set of isopotential compartments. Equation 1 shows the ODE for a single compartment *µ* connected to other compartments *v*, describing the time evolution of membrane potential *V*_*µ*_. Specific membrane capacitance of compartment *µ*, denoted *c*_*µ*_, has units of capacitance per area. The term *i*_*µ*_ has units of electrical current per area and accounts for membrane mechanisms specific to compartment *µ*, such as passive currents and voltage-gated ion channels. External current injection into compartment *µ* is denoted 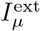, and *A*_*µ*_ is its surface area. The sum in the last term accounts for current contributions from connected compartments *v*, with *g*_*µ,v*_ denoting the conductance per unit area of the connection, which determines the resistive coupling between adjacent compartments and depends on the axial resistivity and the geometry of compartments *µ* and *v*.

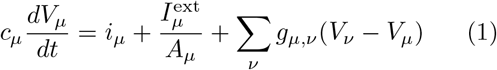

The geometry and connectivity of the cable model strongly influence the cable equations and, thus, the membrane potential produced by the simulator. The goal of `MASCAF` is to provide more realistic cable models for complex morphologies and, therefore, improve the fidelity and quality of multicompartmental simulations.

The `MASCAF` pipeline assumes 3D surface mesh data as input (a triangle mesh of 3D vertices, edges, and faces, e.g., the Wavefront OBJ format (Wavefront Technologies 1990)). For morphologies derived from imaging data, this may require an intermediate membrane-tracing step, typically tracing the neuronal membrane on each slice and converting the result to a triangle mesh. As such, combining `MASCAF` with a membrane-tracing process to produce surface meshes constitutes a complete pipeline from imaging data to multicompartmental simulation. Many software tools exist for semi- or fully automated tracing of cell features in imaging data (Acciai et al. 2016); see Appendix C for a brief description of related software tools.

Cable models typically encode neuronal morphology as an *arborescence*: in graph theory, a directed, rooted, acyclic tree. SWC (Mehta et al. 2023) is a general-purpose cable model format widely used in neuroscience; `MASCAF` cable models are saved in this format. An SWC model describes an arborescence as a list of vertices, each with the following parameters:

- unique index (integer)
- tag (integer)
- spatial coordinates *x, y, z* (float)
- local radius (float)
- parent index (integer)

In a valid SWC model, every non-root node must have exactly one parent, implicitly enforcing acyclicity. `MASCAF` handles cyclic morphologies by breaking cycles at export and writing reconnection directives in the SWC header, which are used to restore loops in the simulator (Section 2.5).

#### Terminology of 3D geometry

Throughout this paper, a *skeleton* denotes a one-dimensional midline representation of the mesh volume. A skeleton may be represented by a structured point set, line set, or 3D graph. A *morphology graph* is a 3D graph whose vertices carry three-dimensional coordinates; it is a downsampled and optimized version of the skeleton that serves as the basis for the cable model. A *cable model* is a 3D graph whose vertices also carry local radius values; it is the discretized morphology used in multicompartmental simulation (and, when represented as a tree, may be written in SWC format). `MASCAF` computes a skeleton from a surface mesh, then constructs the cable model from the mesh and skeleton. Figure 1 (A) shows a simple example of a surface mesh (blue) with an overlaid skeleton (red), and (B) the corresponding cable model with vertices (dark blue), edges (black), and frustum-shaped segments deduced from radii (light blue).

**Fig. 1:**
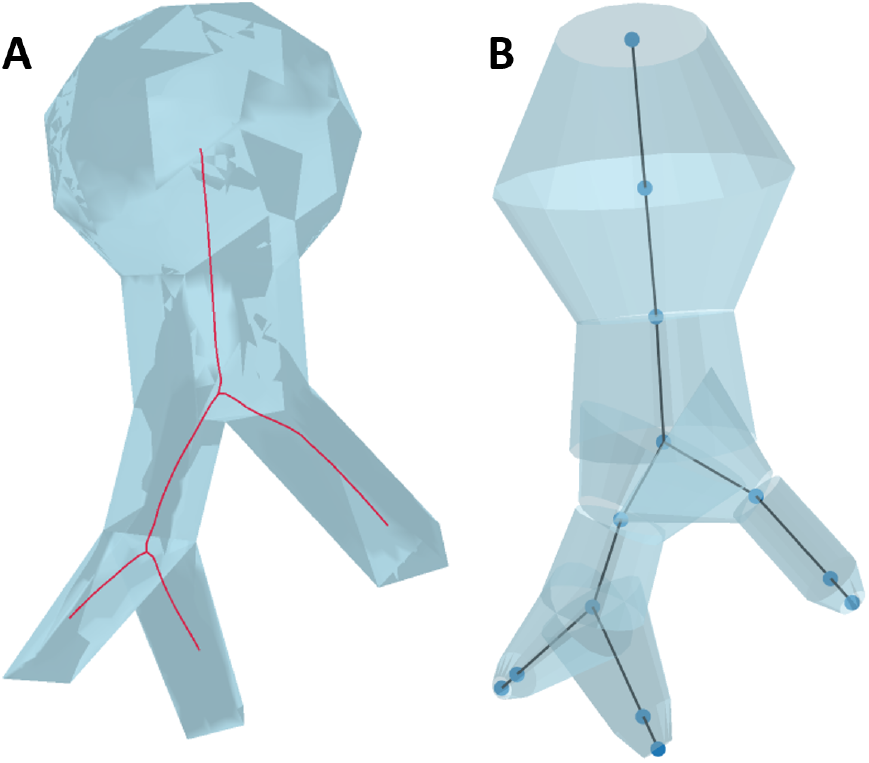
A simple example of a 3D surface mesh with skeleton overlaid in red (A), and the corresponding cable model created using `MASCAF` (B). The cable model shows vertices (dark blue) and edges (black). The surface of the cable model is a union of frustum-shaped segments around each edge, determined by adjacent vertex radii.

### 1.2 Motivation: Toric Spines in the Barn Owl Auditory Pathway

The development of the `MASCAF` pipeline is motivated by the discovery of uniquely shaped den-dritic protrusions, termed toric spines, on space-specific neurons (SSNs) in the barn owl midbrain (Sanculi et al. 2020). SSNs play a central role in the encoding of sound source location (Spezio and Takahashi 2003; Keller and Takahashi 2005; Singheiser et al. 2012); these neurons compute auditory spatial receptive fields via integration of synaptic inputs encoding sound frequency, interaural time difference (ITD), and interaural level difference (ILD) (Peña and Konishi 2001, 2002; Fischer et al. 2009). They are found in the external nucleus of the inferior colliculus (ICx) and arranged topographically to form a map of auditory space (Knudsen and Konishi 1978). The auditory space map in ICx is relayed to the optic tectum (OT) where it aligns and integrates with a visual space map (Knudsen 1982; Knudsen and Brainard 1991). This bimodal map is used to guide orienting movements during nocturnal hunting (Payne 1971; Konishi 1973; Knudsen and Konishi 1979; Moiseff and Konishi 1981; Peña and DeBello 2010).

The ICx is also a major site of experience-dependent plasticity (Knudsen 2002). Young owls raised wearing prismatic spectacles that optically displace the visual field develop adaptive shifts in SSN auditory tuning, which restore bimodal map alignment and behavioral integrity (Knudsen and Knudsen 1989; DeBello et al. 2001; Brainard and Knudsen 1993; Knudsen and Brainard 1991; Knudsen 2002). Manipulations of acoustic experience produce similar effects specific for ITD-frequency combinations (Gold and Knudsen 2000). In both paradigms, auditory tuning in the structure that provides auditory input to ICx, the ICcls, is not altered. Thus, rearrangement of the axonal projections to toric spines and SSNs represents a prime locus for microanatomical plasticity.

Toric spine morphology and connectivity are highly unusual. Compared with typical dendritic spines, toric spines exhibit overall large size, wide necks, a lack of distinct spine heads, and a narrow, tubular geometry (Hering and Sheng 2001; Nimchinsky et al. 2002). Most distinctively, they can exhibit topological holes: tunnels through the membrane that increase the genus of the closed surface mesh. These self-connections represent two distinct ultrastructures: cytoplasmic continuities which must be electrically contiguous, or junctions, which could be electrically contiguous depending on the presence/absence of connexin proteins that form gap junctions. This diversity of electrical properties is likely to powerfully shape the computations performed by SSNs. Highlighting this importance, a single toric spine can receive many synapses from many axons; 49 synapses from as many as 11 axons were observed (Sanculi et al. 2020). Thus, they are structured to act as integrators. A rigorous examination of the relationship between toric spine morphology and synaptic integration is a long-term goal of this work.

On the SSN chosen for high-resolution surface reconstruction from SBEM data, shown in Figure 2, 76 spines were identified, all classified as toric. The toric spines on this cell were observed to have anywhere from zero to nine holes, as well as from one to three necks per spine. Of the 76 toric spines observed, 9 were selected for detailed reconstruction and simulation. These surface meshes are from the reconstruction reported by Sanculi et al. (Sanculi et al. 2020); they are shown in Figure 3 and are publicly available at https://github.com/jmrfox/mascaf/tree/main/data/mesh/processed.

**Fig. 2:**
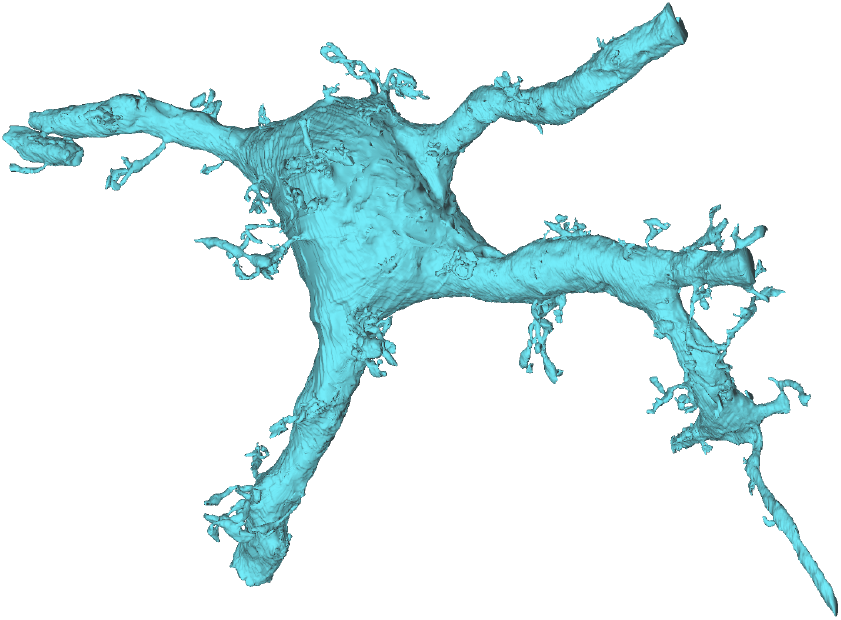
Surface mesh reconstruction of an auditory space-specific neuron in the barn owl midbrain, compiled from electron microscopy data. Large dendritic processes extend out of the EM volume (75 × 45 × 45 *µ*m). Toric spines are visible as thorny structures covering the surfaces of the dendrites and soma.

**Fig. 3:**
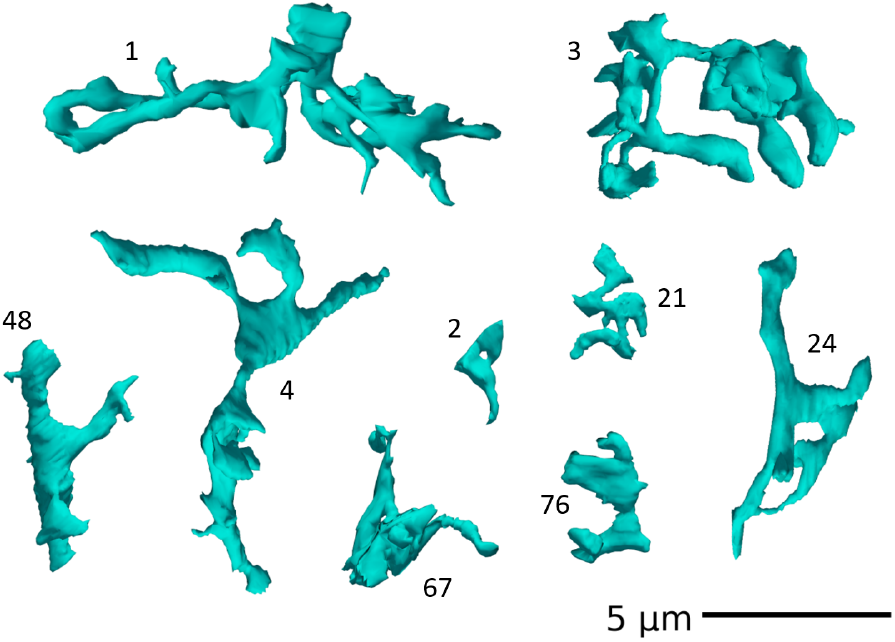
Surface mesh reconstructions of nine selected toric spines from the barn owl ICx neuron shown in Figure 2. Each spine exhibits complex morphology including multiple necks, high curvature, and topological holes.

**Fig. 4:**
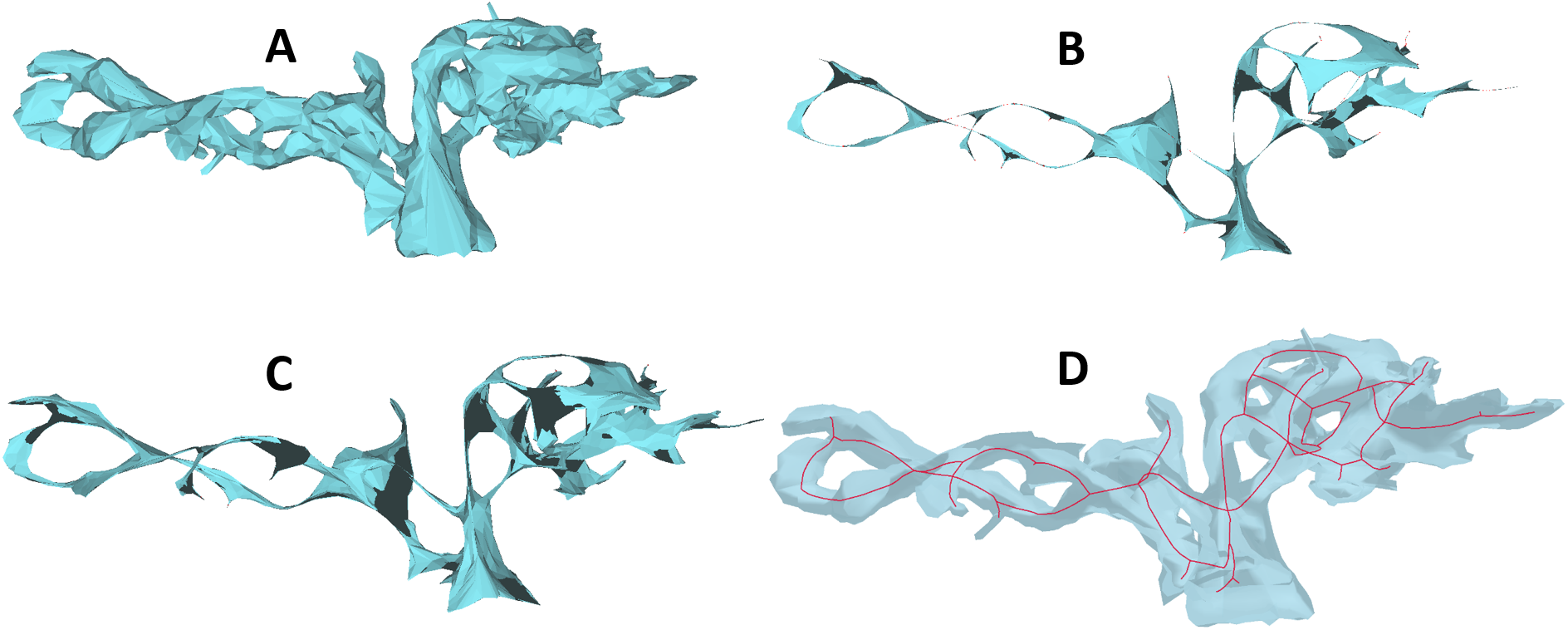
Mean curvature flow skeletonization via `CGAL-TSMS` provides a reliable midline calculation. The original toric spine mesh (A) is iteratively thinned (B, C) to produce a 1-dimensional skeleton (D).

Software tools for neuron modeling often sensibly assume an arborescent structure (Oxford Instruments; Peng et al. 2014; Sato et al. 2000), but the holes of toric spines violate that assumption. The simulators `NEURON` and `Arbor` conveniently allow the establishment of direct self-connections via a gap junction-like mechanism (Carnevale and Hines 2006; Abi Akar et al. 2019), and, therefore, the core modeling challenge is in fitting a valid cable model to the surface mesh. Although development of `MASCAF` was motivated by the challenges presented by toric spine morphologies, which differ from common whole-neuron morphologies in overall scale and geometry, `MASCAF` can be applied similarly to any valid 3D surface mesh, as demonstrated in Figures 1, 9, and 10.

Toward the development of a pipeline for preparing toric spine morphologies for multicompartmental simulation, we identified several key design requirements. The pipeline must robustly compute one-dimensional skeletons from three-dimensional surface meshes, including geometries with high curvature, complex branching, and topological cycles. Structural cycles should generally be retained in the skeleton and the fitted cable model. The cable-model cycle count need not strictly equal the mesh genus, and it may be optimal in some cases to ignore topological holes that do not meaningfully change model structure. The resulting cable model must be exported in SWC or an equivalent format suitable for direct construction of multicompartmental models. To maximize accessibility and reproducibility, the pipeline should operate directly on the surface mesh without requiring additional geometric inputs, rely primarily on deterministic geometric algorithms rather than global optimization or learned models, and be implemented entirely using free and open-source software.

**Table 1:**
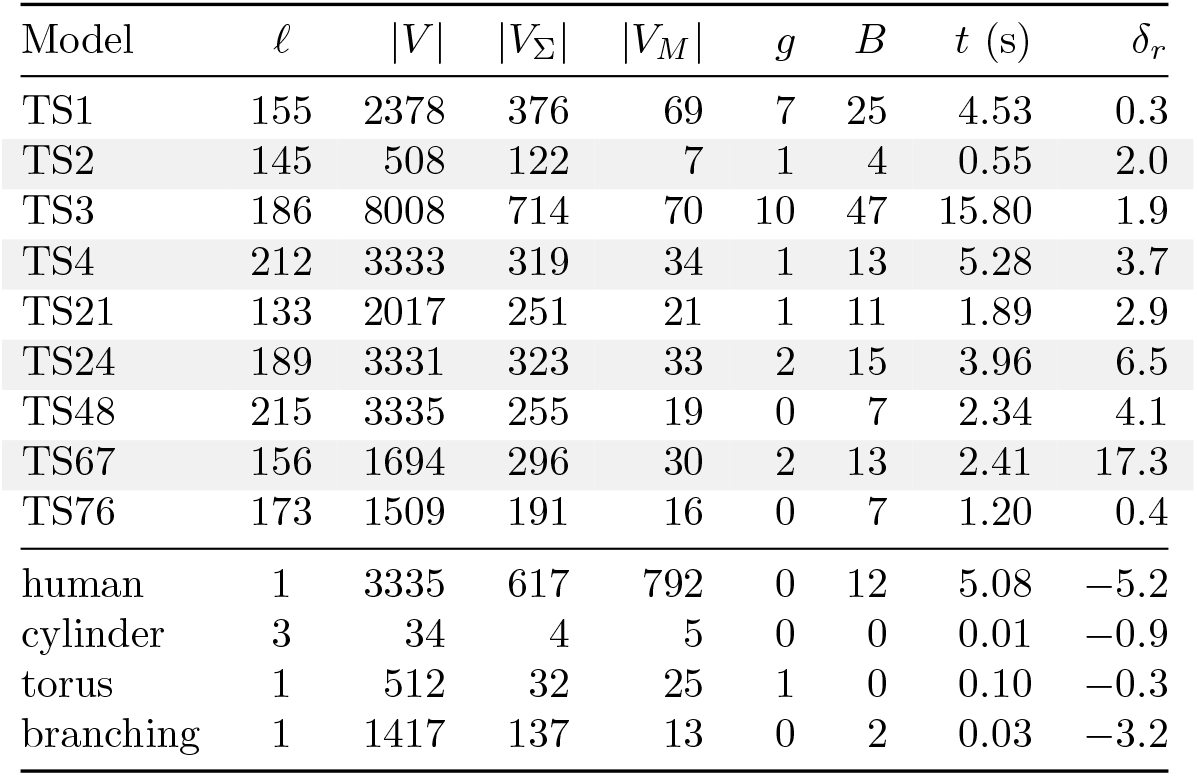
Geometric validation and compute cost. Above the horizontal rule: nine toric spines; those runs used 20 iterations of vertex forcing. Below the rule: a human cortical interneuron (Figure 9) and three synthetic meshes (cylinder and torus in Figure 10, branching model in Figure 1); those runs used no vertex forcing. Terminal extension was used for the branching synthetic mesh and the human neuron. Recorded wall-clock time includes vertex forcing when it is used. *l*: maximum edge length in the cable fitting algorithm. |*V* |: number of vertices in the input surface mesh. |*V*_Σ_|: number of vertices in the skeleton graph *G*_Σ_. |*V*_*M*_ |: number of vertices in the resampled morphology graph *G*_*M*_ after basis optimization. *g*: cyclomatic number of the cable model graph *G*_*M*_ (independent cycles). *B*: number of branch vertices in *G*_Σ_. *t*: end-to-end `MASCAF` wall-clock time in seconds. *δ*_*r*_: the primary geometric error metric, an estimate of radius error in percent, derived from volume and surface area errors.

| Model | ℓ | \|*V*\| | \|*V*<sub>Σ</sub>\| | \|*V*<sub>*M*</sub>\| | *g* | *B* | *t* (s) | *δ*<sub>*r*</sub> |
|---|---|---|---|---|---|---|---|---|
| TS1 | 155 | 2378 | 376 | 69 | 7 | 25 | 4.53 | 0.3 |
| TS2 | 145 | 508 | 122 | 7 | 1 | 4 | 0.55 | 2.0 |
| TS3 | 186 | 8008 | 714 | 70 | 10 | 47 | 15.80 | 1.9 |
| TS4 | 212 | 3333 | 319 | 34 | 1 | 13 | 5.28 | 3.7 |
| TS21 | 133 | 2017 | 251 | 21 | 1 | 11 | 1.89 | 2.9 |
| TS24 | 189 | 3331 | 323 | 33 | 2 | 15 | 3.96 | 6.5 |
| TS48 | 215 | 3335 | 255 | 19 | 0 | 7 | 2.34 | 4.1 |
| TS67 | 156 | 1694 | 296 | 30 | 2 | 13 | 2.41 | 17.3 |
| TS76 | 173 | 1509 | 191 | 16 | 0 | 7 | 1.20 | 0.4 |
| human | 1 | 3335 | 617 | 792 | 0 | 12 | 5.08 | −5.2 |
| cylinder | 3 | 34 | 4 | 5 | 0 | 0 | 0.01 | −0.9 |
| torus | 1 | 512 | 32 | 25 | 1 | 0 | 0.10 | −0.3 |
| branching | 1 | 1417 | 137 | 13 | 0 | 2 | 0.03 | −3.2 |

Although the pipeline emphasizes geometric and topological fidelity, its ultimate purpose is to produce cable models for electrophysiological simulation rather than geometric reconstruction. The cable model serves as the morphological substrate from which multicompartmental simulators derive the electrical properties of the structure. Consequently, throughout this work we prioritize the accuracy of the resulting electrophysiological simulations over purely geometric or visual fidelity.

### 1.3 Contributions

The main contribution of this paper is the Mesh and Skeleton Cable Fitting (`MASCAF`) software (Fox 2026), which fits a cable model to a 3D surface mesh. We explain how the CGAL Triangulated Surface Mesh Skeletonization (`CGAL-TSMS`) algorithm is applied to provide a robust, computationally efficient approach for skeletonizing complex meshes, including those with cycles. Section 2 details methods used for skeletonization, graph optimization, and cable model fitting. In Section 3, we describe validation of our methods in geometric and simulation contexts. Section 4 notes limitations and opportunities for future work. Appendices provide a pipeline flowchart (A), algorithms (B), and a brief description of related software tools (C).

By enabling the conversion of high-resolution mesh reconstructions into multicompartmental simulation models without reliance on proprietary software, this pipeline lowers technical and financial barriers to morphology-driven computational neuroscience.

## 2 Methods

This section describes steps of the `MASCAF` pipeline for fitting a cable model to a 3D surface mesh. A flowchart of the `MASCAF` pipeline is shown in Appendix A. Mesh operations described in this section use the `trimesh` Python package (Dawson-Haggerty et al. 2019), and graph operations use `networkx` (Hagberg et al. 2008).

### 2.1 Input Data & Mesh Preprocessing

Mesh data is eligible for the `CGAL-TSMS` algorithm if and only if it is a single component, has no self-intersections and no boundaries (i.e., is “watertight”). If any of these prerequisites are not met, the operation will return an error before running. Meshes derived from imaging or contour data are made watertight either by standard mesh hole-filling routines or by alpha-wrapping before skeletonization (Alliez et al. 2026).

The `MASCAF` Python code offers two mesh pre-processing operations using `CGAL`: repair (Ohori 2026), and simplification (Cacciola et al. 2026). If the original mesh has a higher resolution than is needed for skeletonization, one may apply the simplification operation to reduce the number of vertices and edges. This will speed up the subsequent skeletonization operation, which scales with the number of mesh vertices. In our case, some meshes converted directly from contour data had several hundred thousand edges, and simplifying them to a few thousand edges greatly sped up operations without sacrificing fidelity.

### 2.2 Mean Curvature Flow Skeletonization via CGAL-TSMS

The central algorithm in `CGAL-TSMS` is mean-curvature flow skeletonization (MCFS), which iteratively displaces vertices according to local curvature until the mesh collapses to a one-dimensional skeleton (a curve for genus-0 meshes, or a graph that may contain cycles). `CGAL-TSMS` can be applied within the `MASCAF` Python code, in the `CGALLab` program (GUI), or programmatically in C++ code using the `CGAL` library (The CGAL Project 2026; Gao et al. 2026). A thorough description and analysis of MCFS can be found in the original paper (Tagliasacchi et al. 2012). A brief and application-focused description of the MCFS process is as follows.

#### 2.2.1 Mathematical Foundation

##### Mean curvature flow skeletonization

(MCFS) is a dynamical process applied to the closed 2D surface that contracts it toward a 1D midline. Consider a map **p** : *U*⊂ R^2^ → R^3^ where **p**(*u, v*) = (*x*(*u, v*), *y*(*u, v*), *z*(*u, v*)) and *U* is a closed domain in the *u, v* plane. We denote our closed 2D surface *S* ≡ {**p**(*u, v*) | *u, v* ∈ *U}*, which encloses a 3D volume Ω, so *S* = *∂*Ω. Under MCFS, the time rate of change of the surface is equal to the local inward normal −**n** (where **n** is the outward unit normal, so −**n** points *into* the volume Ω) scaled by the local mean curvature *H* = (*k*_1_ + *k*_2_)*/*2, where *k*_1_ and *k*_2_ are the principal curvatures.

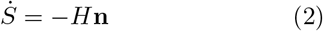

In practice, however, the vector *H***n** is challenging to compute. Instead, MCFS invokes the Laplace-Beltrami operator on *S*, Δ_*S*_, which is a generalization of the standard Laplacian operator to curved subspaces.

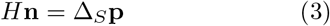

The skeleton solution **p**^*^ is interpreted as having infinitesimal volume and surface area, and so mean curvature flow halts at that point.

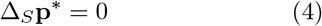

#### 2.2.2 Discretized Mean Curvature Flow Skeletonization

In the discrete representation, *S* is approximated by a 2D surface mesh *M*_*S*_, which is defined by vertices, edges, and triangular faces. Similarly, we denote the internal volume *M*_Ω_. The mesh has vertices *V* = [**v**_1_, **v**_2_, …, **v**_*n*_], the locations of which constitute a discrete approximation of the function **p**. The `CGAL-TSMS` algorithm updates vertex positions with an implicit Euler step, *V* ^*t*^ *V*→ ^*t*+1^ until total change in *V* is below a convergence threshold. Further details about the solution process are discussed in the original publication (Tagliasacchi et al. 2012).

To understand the parameterization of MCFS relevant to `MASCAF`, we can consider the total “energy” for a configuration *V* at time *t*,

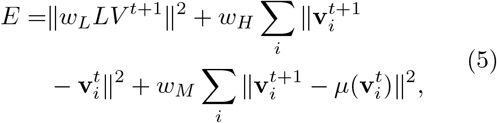

which is consequently minimized during MCFS. Here, *L* is the discrete Laplace-Beltrami operator, *w*_*L*_, *w*_*H*_, and *w*_*M*_ are weight parameters, and *µ*(·) is a map from a vertex to its nearest Voronoi pole. Voronoi poles constitute a sparse approximation of the mesh’s midline in the form of a point cloud; they are useful here in guiding *V* toward the midline, but do not constitute a solution to MCFS on their own. The three weights are constrained by a partition-of-unity property; only *w*_*H*_ and *w*_*M*_ must be set by the user, and *w*_*L*_ is computed internally. For each term in Equation 5, a larger corresponding weight value emphasizes that term’s influence in the solution: a larger *w*_*H*_ leads to smaller changes per time step, and a larger *w*_*M*_ produces vertices closer to Voronoi poles.

The `CGAL-TSMS` C++ implementation has the following exposed parameters.

- `is medially centered` (boolean): Is *w*_*M*_ *>* 0?
- `quality speed tradeoff` (float): *w*_*H*_
- `medially centered speed tradeoff` (float): *w*_*M*_

In `MASCAF`, `is medially centered` will generally be `true` because *w*_*M*_ *>* 0 provides a solution with vertices nearer to the mesh’s midline. The values of *w*_*H*_ and *w*_*M*_ affect skeleton quality, smoothness, branching density, and compute cost, and we cannot provide a single parameterization that guarantees high-quality skeletons in general. Optimal balancing of parameters ultimately depends on features of the input data. For example, larger *w*_*H*_ can simultaneously increase skeleton detail and branching density, so an optimal value results in a balance of those effects. On the other hand, fitting of significant topological cycles is observed to be very robust under reasonable parameterizations. Suboptimal skeleton results can be optimized within `MASCAF` using methods discussed in Sections 2.3.2 and 2.3.3. For all our toric spine meshes, skeletons were generated with *w*_*H*_ = 0.5 and *w*_*M*_ = 5.

#### 2.2.3 Skeleton Data: Polylines and the Skeleton Graph

The skeleton Σ computed by `CGAL-TSMS` is specified in standard *polylines* format and consists of a set of *m* ordered lists of 3D points:

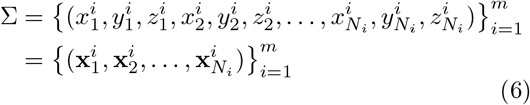

where 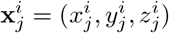 is a single 3D point (the *j*-th point in polyline *i*).

The locations of branch points (degree 3 or more) and terminal points (degree 1) can be deduced from polyline data, which allows us to construct the *skeleton graph G*_Σ_. Each of the first and last coordinates in a polyline, 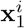 and 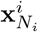, must be either a branch point or a terminal point. If the coordinate of an endpoint matches an endpoint of a different polyline within a small tolerance *ε* (e.g., 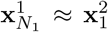), then the two poly-lines are connected at that point, and so it marks a branch point. If an endpoint is unique, then it is a terminal point. Points with *j* = 2, 3, …, *N*_*i*_ − 1 must have degree 2. Furthermore, because *M*_*S*_ is a single component, *G*_Σ_ must be a single component. This is sufficient information to construct the skeleton graph, which is an undirected graph *G*_Σ_ with vertices *V*_Σ_, and edges describing the connectivity structure implicit in the ordering of Σ.

#### 2.2.4 Skeleton Quality

For a general three-dimensional surface mesh, there is no unique optimal one-dimensional skeleton. The medial axis transform of a 3D enclosed volume is generally a composite of 2D sheets, 1D curves, and 0D junctions (Blum 1967; Siddiqi and Pizer 2008). Skeleton quality must instead be judged with respect to the intended application. In `MASCAF`, a suitable skeleton should traverse the full extent of the enclosed volume, remain near its geometric midline, preserve the relevant topology, and contain sufficient branches to represent the morphology without introducing unnecessary geometric complexity. The broader implications of this non-uniqueness are discussed in Section 4.1.

The skeleton produced by `CGAL-TSMS` may not satisfy all of these criteria for every parameterization. While choosing *w*_*M*_ *>* 0 encourages contraction toward the medial region of the mesh, individual skeleton points may remain displaced from the local midline, and in some cases may lie outside the closed surface. Small deviations are generally tolerated by the subsequent radius-fitting procedure. However, when a skeleton vertex lies entirely outside the enclosed volume, a meaningful local radius may not be recoverable, and fitting at that location can fail.

Branching density presents a separate challenge. A skeleton with too few branches may fail to represent distinct morphological processes, whereas excessive or spurious branches may over-fill the original volume once radii are assigned. The latter error cannot be assessed completely from the unweighted skeleton alone, because its geometric effect depends on the fitted radii and on the resulting union of cable segments. After radius fitting, branching quality can therefore be evaluated through volumetric reconstruction measures, including overlap between the reconstructed cable volume and the original mesh volume. Before radii are available, branching density must be estimated from the spatial relationship between the skeleton and the mesh surface. `MASCAF` provides a graph-pruning operation that can remove short or otherwise spurious branches before the final cable model is constructed.

When the initial skeleton is insufficiently centered or contains excessive branching, the user may modify *w*_*H*_ and *w*_*M*_ and rerun `CGAL-TSMS`. In practice, these objectives can compete. Increasing *w*_*H*_ generally preserves more of the local surface geometry and can improve morphological coverage, but it may also increase the number of branches retained during contraction. Conversely, stronger contraction toward the medial region may simplify the skeleton while reducing its ability to follow fine morphological features. Consequently, it may be difficult to identify a single parameterization that produces both a well-centered skeleton and an appropriate branching density. In the following section, we introduce an optimization procedure that refines the positions of morphology-graph vertices after the initial graph basis has been constructed from the skeleton.

### 2.3 Cable Model Fitting

Construction of the cable model proceeds in stages: constructing the base morphology graph *G*_*M*_ from the skeleton graph *G*_Σ_, optional basis optimization (pruning, snapping, and forcing), fitting local radii by measuring local distances to the mesh surface, and global surface-area normalization. A pseudocode description of the fitting algorithm is included in Appendix B.2.

#### 2.3.1 Basis Construction: the Morphology Graph

The morphology graph *G*_*M*_ (*V*_*M*_, *E*_*M*_) is an undirected graph that will form the spatial and structural basis for the final cable model. The vertices *V*_*M*_ are instantiated with position values from the skeleton graph *G*_Σ_. Typical skeletons from `CGAL-TSMS` have a higher spatial density of points than is useful for a cable model, so we resample *G*_Σ_ when computing the initial *G*_*M*_. We first identify a set of all terminal and branch vertices as *C*(*V*_Σ_) = {**v** ∈ *V*_Σ_ | deg(**v**) ≠ 2}, then tabulate the set of *N* unique non-branching sections, 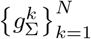. A non-branching section is a maximal edge-connected subgraph of one of two types: (*i*) a path whose endpoints lie in *C*(*V*_Σ_) and whose interior vertices all have degree 2, or (*ii*) a cycle whose vertices all have degree 2. The skeleton graph *G*_Σ_ is decomposed into non-branching sections 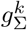 by tracing all maximal paths emanating from vertices in *C*(*V*_Σ_) and then tracing any remaining cycles. This yields an edge-disjoint decomposition of the graph into maximal non-branching sections: 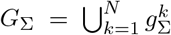. Given a global maximum edge length parameter *l*, each 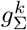 with arc-length 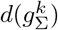 is resampled to have 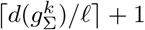 evenly spaced vertices, including the endpoints, to give a new non-branching section *g*^*k*^. Finally, the morphology graph is instantiated as the union of all of the resampled sections, 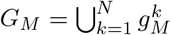. A section shorter than the maximum edge length *l* becomes a single edge. The morphology graph does not keep more than one edge between the same two vertices, so a cycle that then passes through only two vertices disappears. `MASCAF` warns when this downsampling loses a cycle, and the fit is still allowed. The user controls that choice with *l*. Dropping the cycle can be useful when `CGAL-TSMS` produces a small cycle that should not be kept. This gives a robust process to compute basis nodes from the skeleton while maintaining global control over edge length simply via *l*.

Basis optimization, optional refinement of *V*_*M*_ positions to improve the quality of the final cable model, proceeds in three phases: pruning of short terminal branches, vertex snapping, and vertex forcing. The latter two phases are described in the following two sections, and a pseudocode description of the basis optimization algorithm is given in Appendix B.1. Since vertex snapping and forcing scale with the number of vertices targeted, we intentionally apply these operations after the resampling *G*_Σ_ → *G*_*M*_.

#### 2.3.2 Basis Correction: Vertex Snapping

Although rare, Σ will sometimes contain points fully outside *M*_*S*_. This may occur when using sub-optimal parameterizations of `CGAL-TSMS`, and is more likely to occur near regions of high curvature. This can cause errors when fitting radii with `MASCAF`. Vertex snapping identifies vertices outside the mesh volume and moves them inside by updating their positions. Figure 5 illustrates the vertex snapping process.

**Fig. 5:**
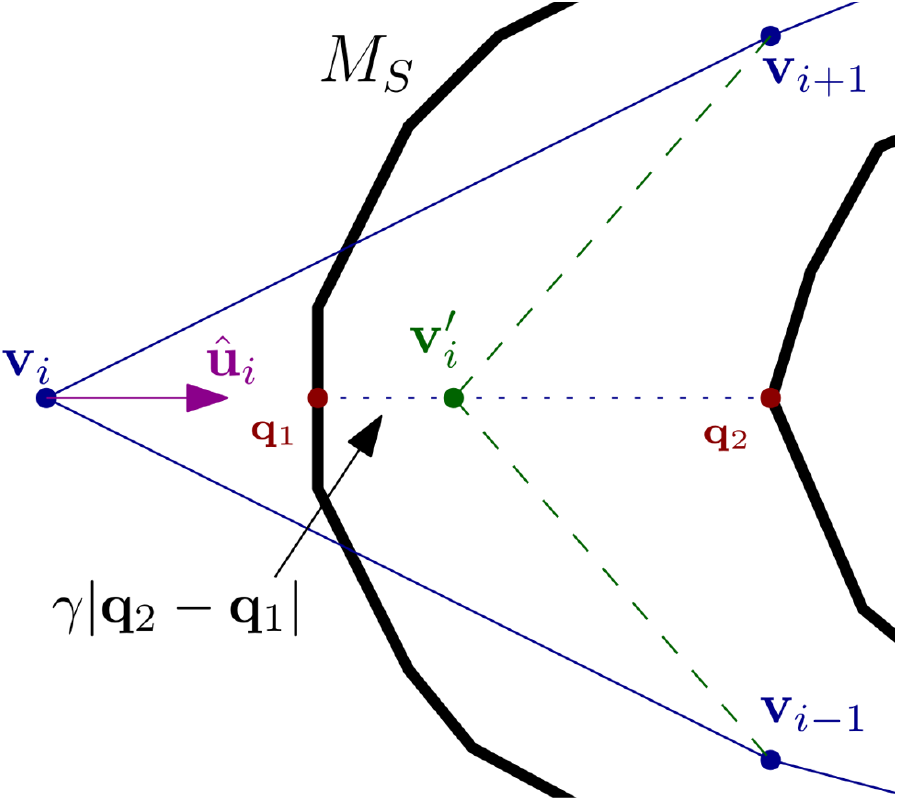
Vertex snapping for a vertex **v**_*i*_ (blue) outside the mesh surface *M*_*S*_ (black). A ray in the direction of the unit vector **û**_*i*_ (magenta) intersects the surface at ordered points **q**_1_ and **q**_2_ (red). The updated vertex 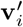 (green) lies on the chord between them at fraction *γ* of the distance from **q**_1_ toward **q**_2_.

Let *σ*(*M*_*S*_, **v**_*i*_) be the point on *M*_*S*_ nearest to **v**_*i*_, and define the unit direction

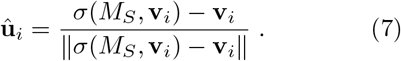

Casting a ray from **v**_*i*_ along **û**_*i*_ yields ordered intersection points {**q**_*j*_} with faces of *M*_*S*_. The snapped position is a fraction *γ* of the chord between the first two intersections,

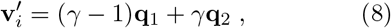

where *γ* controls the placement along the chord (*γ* ≈ 1*/*4), provided that the chord length ∥**q**_2_ − **q**_1_∥ exceeds a small threshold relative to the mesh size.

Although using *γ* = 1*/*2 would very often place 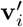 nearer to the mesh midline, this can also lead to extreme angles between the edges of *G*_*M*_ adjacent to 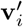, thereby introducing artificial discrepancies between the tangents of *G*_*M*_ and those of *M*_*S*_. This procedure is still inexpensive when only a few vertices lie outside the mesh, and is often sufficient to proceed with radius fitting. If higher midline accuracy is necessary, we continue to the vertex forcing phase of basis optimization.

#### 2.3.3 Basis Refinement: Vertex Forcing

Vertex forcing uses an iterative method to move any vertices that are inside the mesh closer to the midline. Figure 6 illustrates one iteration. Terminal vertices are typically held fixed (and optionally branch vertices), so that connectivity anchors are preserved while non-terminal vertices are refined. This process moves a vertex **v**_*i*_ along an iterative trajectory described by 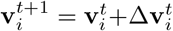. The total force is a mixture of two contributions,

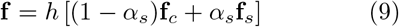

where **f**_*c*_ is a centering force, **f**_*s*_ is a smoothing force, *h* ≥ 0 controls the step scale, and *α*_*s*_ controls the smoothing fraction. To prevent a step from carrying a vertex outside the mesh, we limit the possible change at each iteration. We define the maximum step distance *f*_max_ to be a tunable constant fraction (default 1*/*2) of the distance from 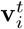 to the nearest face of *M*_*S*_ in the direction of **f**. The applied step is then

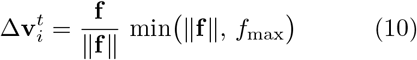

**Fig. 6:**
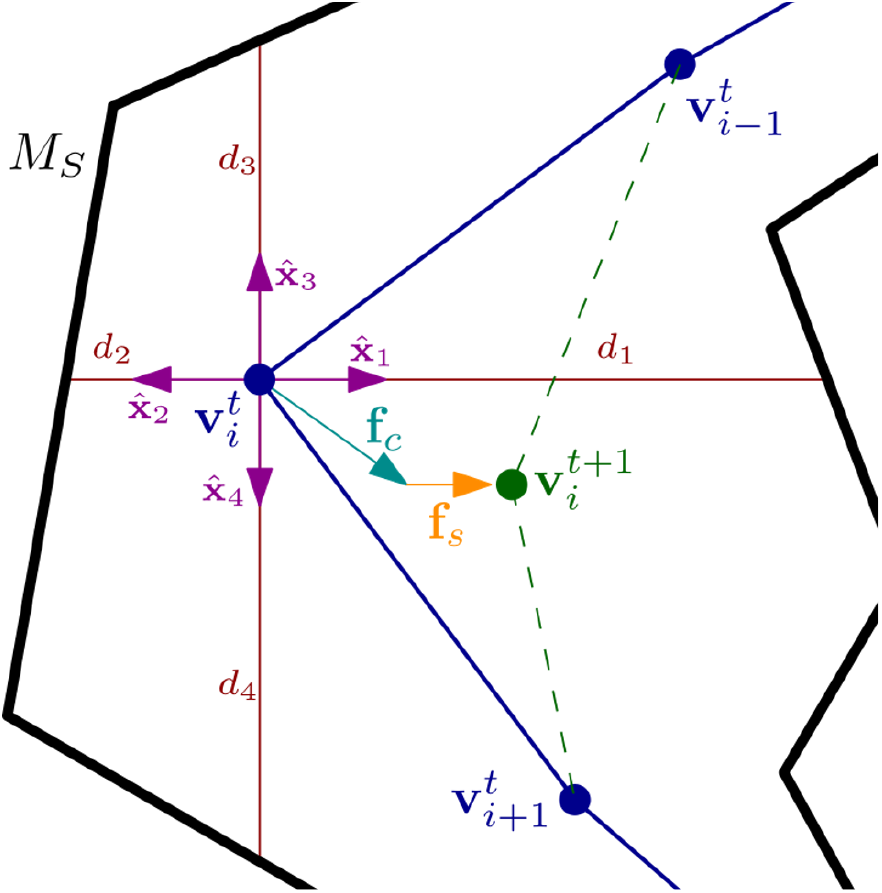
One vertex-forcing iteration for an interior vertex 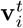 (blue), projected onto a 2D plane. Distances *d*_*j*_ (red lines) from 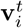 to the mesh *M*_*S*_ (black) along four ray directions given by unit vectors 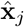 (magenta) determine the centering force **f**_*c*_ (cyan) according to Equations 11 and 12. The smoothing force **f**_*s*_ (orange) pulls **v**_*i*_ toward neighboring vertices. Note that the step scale *h* and smoothing fraction *α*_*s*_ are omitted from this figure for clarity. Vector directions and magnitudes are chosen purely for illustration.

**Fig. 7:**
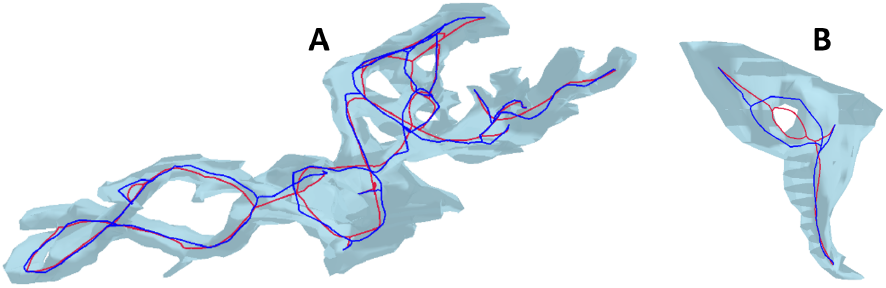
Two toric spine meshes rendered with downsampled skeletons and optimized morphology basis graphs. Skeleton-graph vertices from `CGAL-TSMS` are shown in red, and morphologygraph vertices after `MASCAF` basis optimization are shown in dark blue. Note that, in (B), the initial skeleton is not fully contained within the mesh, while the optimized morphology graph is.

The purpose of the centering force **f**_*c*_ is to push the vertex away from the surface using ray-tracing. To compute the centering force on 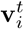, we cast *N*_*R*_ rays from the vertex with unit directions 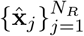. By default, *N*_*R*_ = 6 with unit vectors aligned with the positive and negative Cartesian axes; denser sampling on the sphere (e.g., Fibonacci sampling) can be used when higher accuracy is desired, at greater computational cost. Along each ray, we compute the scalar distance *d*_*j*_ measured from 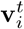 to the nearest intersection with a face of *M*_*S*_. The centering force **f**_*c*_ is the weighted sum of the ray unit vectors,

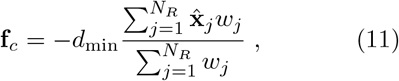

kept at its weighted magnitude (not re-normalized to unit length), where the weight for direction 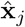 is

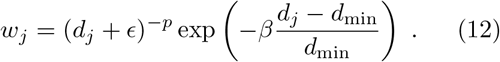

At each iteration, we compute 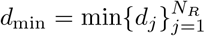, the minimum distance from the vertex to *M*_*S*_ along the rays, which is then used to scale the force weights.

A weight *w*_*j*_ has two multiplicative contributions, one repulsive, and one localizing. The repulsive factor is (*d*_*j*_ + *ϵ*)^*−p*^, where *ϵ* is a small number to avoid division by zero, and *p* is a parameter controlling the shape of the repulsion. This factor is larger for small *d*_*j*_, and the nonlinearity introduced by the exponent *p* (typically *p*∈ [1, 2]) allows for small distance repulsions to be further emphasized over large ones. The exponential factor is for localizing; it allows for the centering force to be dominated by short-range repulsion, and effectively ignores long-range effects. The localizing term exponentially downweights contributions with *d*_*j*_ *> d*_min_. The parameter *β* controls the shape of localization, and is similar in effect to *p*. This weight design produces reliable local centering and provides the user with global control. Default values in `MASCAF` of *p* = 1 and *β* = 2 are adequate for general applications. Note that each term may be turned off by setting its corresponding parameter to zero.

The smoothing force **f**_*s*_, introduced to suppress jitter and maintain spatial coherence, points the vertex 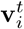 toward the average of positions in the set of neighboring vertices 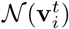:

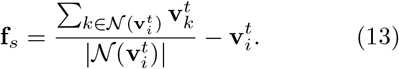

While **f**_*c*_ approaches zero when 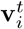 is centered (neglecting long-range forces), **f**_*s*_ does not. Therefore, the practical use of vertex forcing is not to run until ∥**f**∥ ^2^ goes to zero, but rather to either run until ∥**f**_*c*_∥ ^2^ crosses below a threshold, or simply for a fixed number of iterations *T*. In practical applications of vertex forcing, we find that relatively small iteration counts (*T <* 50) produce satisfactory refinement.

#### 2.3.4 Computing Radii

The local radius of each vertex in *V*_*M*_ is computed via the following process. Consider a target vertex **v**_*i*_ ∈ *V*_*M*_. For each adjacent edge *e*_*ij*_, we define a cross-section plane through **v**_*i*_ with normal **e**_*ij*_. Where the plane intersects the mesh, a polygonal cross-section with area *A* is computed using standard mesh manipulation algorithms (`trimesh` (Dawson-Haggerty et al. 2019)). We evaluated several different radius calculation strategies, including comparisons of surface-area-preserving and volume-preserving approaches. Although surface area plays a significant role in the cable equations, we found that a surface-area-preserving approach to radii calculations resulted in noisy radius distributions and larger geometric errors. The most accurate and practically robust strategy we found calculates the radius of a disk with the same area as the cross-sectional polygon; thus 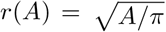. However, we acknowledge that surface area preservation may be an optimal strategy in some applications, and for that reason the option is included in the `MASCAF` code. This process is repeated for each edge *e*_*ij*_ connected to **v**_*i*_, producing a set of radii 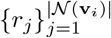. Finally, the assigned radius *r*_*i*_ is a reduction of those, typically mean (the default) or median. Once every vertex in *V*_*M*_ has a radius, we treat *G*_*M*_ together with its frustum segment geometry as the cable model.

### 2.4 Fine Tuning: Surface Area Normalization & Terminal Extension

Compartment surface area has a significant effect on electrical simulations (Equation 1), augmenting or suppressing the effects of external currents. `MASCAF` therefore applies a final normalization step: a global scale multiplies all radii so that the sum of segment lateral surface areas matches the mesh surface area. Branch-point overlap, which motivates that scale, is estimated only when computing geometric errors (Section 3.1.1). Multicompartmental simulators typically use lateral membrane area in the cable equations, so normalization by default excludes terminal end caps (Carnevale and Hines 2006). Relative scales between radii within the same model are preserved, and thus relative compartment membrane areas are preserved under uniform specific membrane properties.

Lastly, `MASCAF` includes a convenient method to extend all terminal points (degree-1 vertices) of a cable model with an additional segment, scaled to the local radius. Due to medial centering, skeletons computed with `CGAL-TSMS` often feature branches that stop short of the end of the mesh. Depending on mesh geometry, this may lead to substantial geometric errors. For example, note how the skeleton of the branching geometry shown in Figure 1 (A) stops near the center of the bulk, but the cable model (B) has an additional terminal point at the top (and also ones on the lower branch terminals), added using terminal extension. The terminal-extension functionality adds new terminal points so the cable model extends beyond the skeleton, leading to reduced geometric errors, though it is not necessarily appropriate for all cases.

### 2.5 Simulator Integration

Since the SWC format is limited to describing acyclic graphs, cycles in the cable model must be resolved prior to export. `MASCAF` addresses this limitation by effectively breaking graph cycles and writing reconnection directives to the SWC file header. On each detected cycle 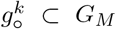, one degree-2 vertex **v** is selected. This vertex is duplicated to create a clone **v**^*′*^ with identical attributes, and one of the edges terminating on **v** is updated to instead terminate on **v**^*′*^. The two duplicate vertices are distinct in the SWC specification and are not connected by an edge. This operation thus breaks the cycle while preserving geometric embedding. `MASCAF` adds metadata describing each broken cycle to the SWC file header, including the indices of **v** and **v**^*′*^.

After initializing the morphology in simulation software, the original loop 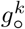can be restored by adding a direct connection between **v** and **v**’. In `Arbor`, for example, existing gap-junction functionality (normally used to connect two neurons) can be repurposed to place a direct connection between the corresponding segments, exposing a conductance parameter for each connection (Abi Akar et al. 2019). A similar procedure can be implemented in `NEURON` (Carnevale and Hines 2006). We set the conductance of this connection (in siemens) to the reciprocal of axial resistance as a lumped electrical approximation of full cytoplasmic continuity (Carnevale and Hines 2006). As such, the `MASCAF` approach allows the modeler to recreate topological cycles at the simulation level while still using the common SWC format.

## 3 Results

This section presents selected results from `MASCAF` and covers geometric and simulation validation of the fitted cable models. As a representative example, we report validation metrics for toric spine #1 (TS1), fitted from a mesh with 2378 vertices. The original mesh, skeleton, and two cable model results for TS1 are shown in Figure 8. Before cycle-breaking, the cable model had 69 vertices, 75 edges, and 7 independent cycles; after cycle-breaking and SWC export, the arborescence has 76 vertices and 75 edges. Other attributes and error estimates are given in Table 1.

**Fig. 8:**
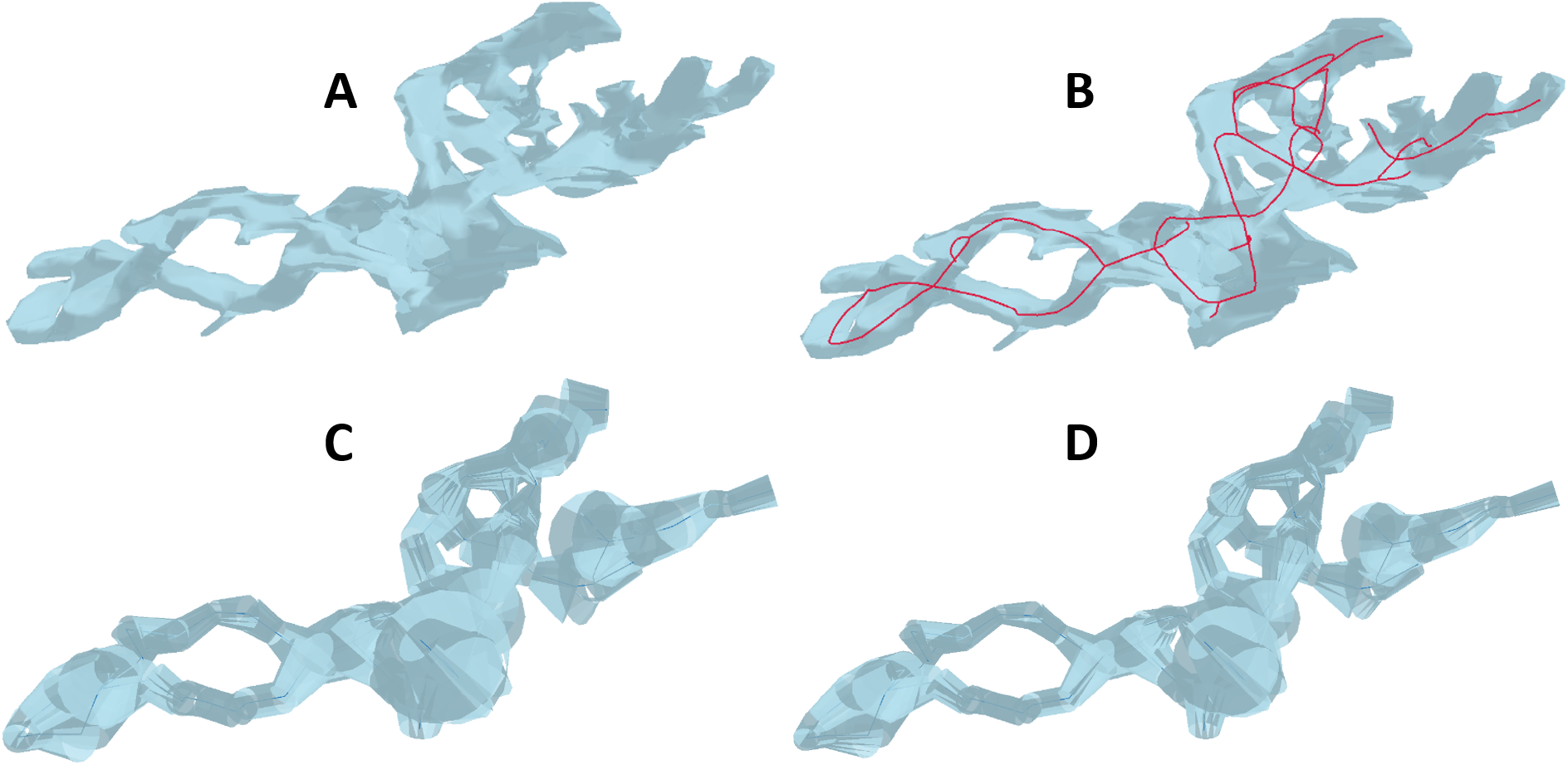
Toric spine #1. (A) Original surface mesh. (B) Mesh (blue) with the skeleton (red). (C) Fitted cable model. (D) Final cable model after normalizing surface area to match that of the mesh. The fitted model (C) and area-scaled model (D) are different but visually quite similar, which suggests the radius calculation of Section 2.3.4 is accurate. Error estimates are listed in Table 1.

To illustrate the applicability of `MASCAF` to full neuronal morphology data, we include results for a human neuron, shown in Figure 9. The morphology mesh data for this human cortical interneuron (middle temporal gyrus), reconstructed by the Allen Institute, was provided by NeuroMorpho.org (Koch and Jones 2016; Tecuatl et al. 2024; Ascoli et al. 2007).

**Fig. 9:**
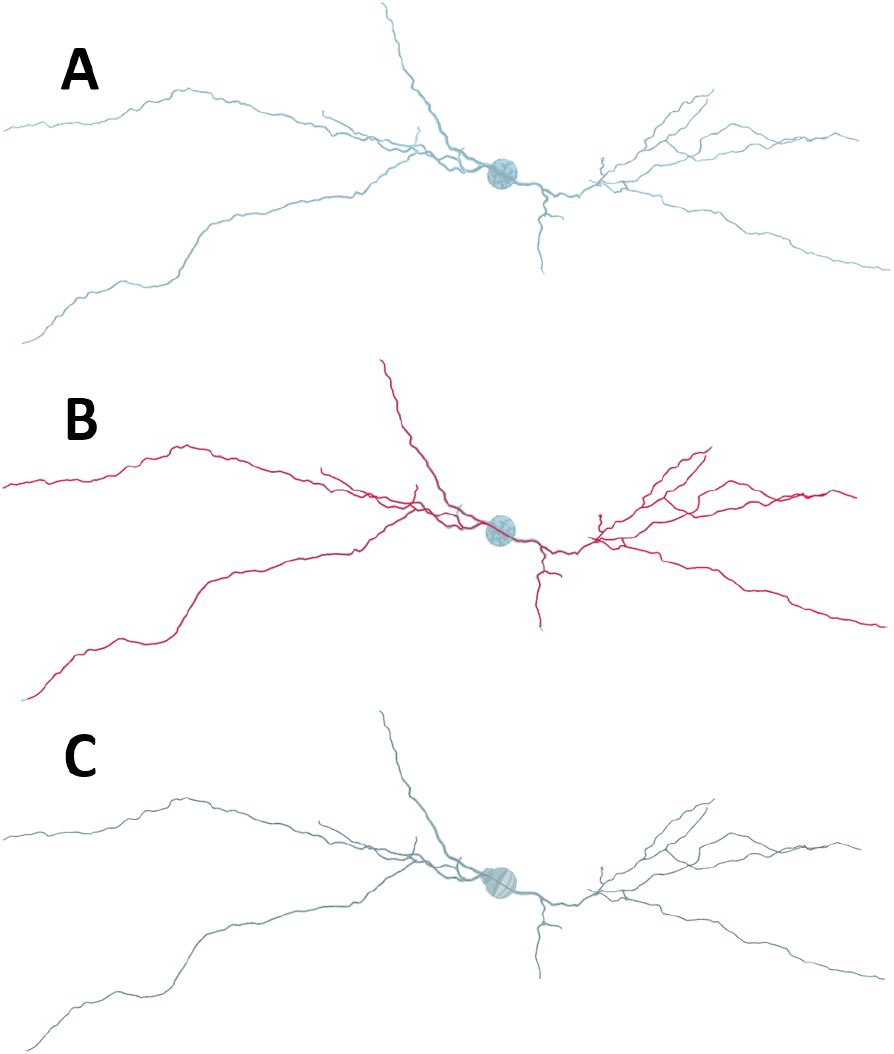
Three steps illustrating `MASCAF` applied to the surface mesh of a human cortical interneuron from the middle temporal gyrus. (A) shows the neuron mesh, (B) shows the mesh (blue) with the skeleton (red), and (C) shows the cable model. The fitted cable model is similar to the original mesh. Geometric errors for this example are listed in Table 1.

### 3.1 Geometric Validation

To evaluate geometric accuracy of results, we construct a frustum-based geometric proxy from the fitted cable model and compare its volume and lateral surface area to those of the original mesh *M*_*S*_. Figure 10 shows `MASCAF` results for two synthetic meshes with tractable geometry: a straight cylinder and a torus. Consider the cable model graph *G*_*M*_ (*V*_*M*_, *E*_*M*_) with edge lengths and vertex radii from the fitting pipeline. For each pair of vertices **v**_*i*_, **v**_*j*_ ∈ *V*_*M*_ that share an edge *e*_*ij*_ with length |**e**_*ij*_|, we construct a *segment* geometry, a frustum with radii *r*_*i*_, *r*_*j*_, which has volume

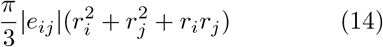

and lateral surface area (not including end caps)

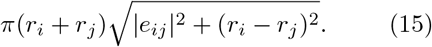

**Fig. 10:**
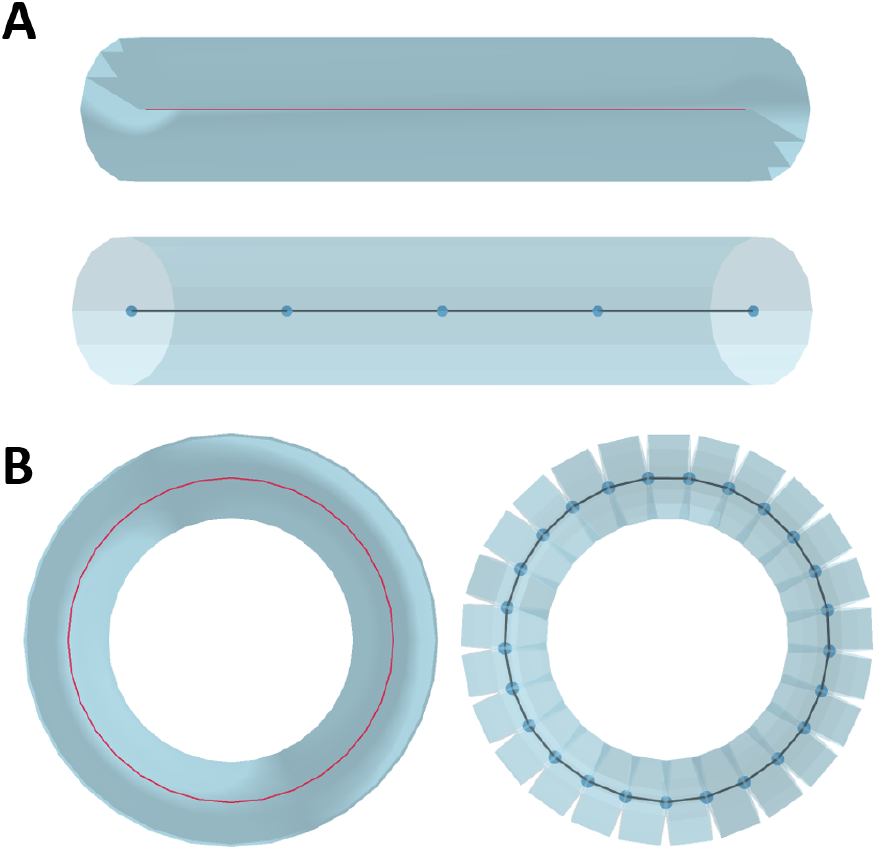
Known-shape validation on synthetic meshes. (A) Cylinder: input surface mesh with skeleton (top) and fitted cable model (bottom). The cable model has 5 vertices and 4 compartments. Segment boundaries are hard to see because the segments are parallel and meet. (B) Torus: input surface mesh with skeleton (left) and fitted cable model (right). This cable model illustrates how the overlap of segments around a degree-2 vertex (see Section 3.1.1) tends to cancel with the opposing gap.

The total volume of the cable model is estimated as the sum of all segment volumes. The total surface area used for comparison with the mesh is the sum of all segment lateral surface areas.

The `MASCAF` Python code includes a geometric validation system that compares the surface area and volume of the mesh to those of the cable model, with and without overlap subtraction (see Section 3.1.1). Mesh genus counts every hole. Very small holes that do not substantially change the morphology can be dropped when the mesh is simplified to a skeleton, and resampling can remove a short cycle, as described in Section 2.3.1. The cable model can therefore have fewer cycles than the genus. For toric spine #1, the mesh genus measured with `trimesh` is 9, but two of those holes are very small pinch points rather than structural tunnels, and the fitted cable model has 7 independent cycles. The `MASCAF` validation routine reports the measured mesh genus and the cable-model cycle count so they can be compared.

Table 1 reports geometric validation metrics and pipeline cost for the nine toric spines used in this study and for four other meshes: a human cortical interneuron and the synthetic cylinder, torus, and branching models. The toric-spine runs used 20 iterations of vertex forcing. The human neuron and the synthetic meshes used no vertex forcing because their skeletons were already accurate. Because only the toric-spine runs include vertex forcing, times should not be compared across the two blocks as if the forcing setting were the same. End-to-end wall-clock time was measured on a single workstation (Intel Core i9 CPU at 3.60 GHz, 8 cores) for the fitting runs underlying the table. All listed runs required peak RAM between 151.8 MB and 231.1 MB. Overall run time is governed mainly by skeleton size |*V*|_Σ_, and vertex forcing is the most costly phase. `CGAL-TSMS` scales appreciably with the mesh vertex count |*V*|; elsewhere in `MASCAF`, |*V*| enters mainly because the input surface must be retained in memory for geometric queries, whereas the cable fitting algorithm itself is largely independent of |*V*|. For analysis of the time demands of `CGAL-TSMS`, we refer readers to Tagliasacchi et al. (2012).

The values in Table 1 are from automated runs, with vertex forcing as described above. Lower errors are possible with more careful case-by-case tuning. The *δ*_*r*_ column is the primary geometric error metric: an estimate of radius error in percent, derived from volume and surface area errors,

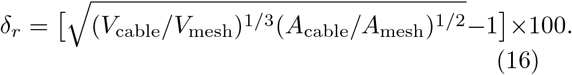

#### 3.1.1 Overlap estimation

A fundamental problem in geometric fitting of cable models arises from volumetric and surface-area overlap at vertices with more than two edges (branch nodes). At each vertex of degree greater than 2, there is a systematic positive overlap in volume and surface area of the connected segments. Depending on attributes of the morphology like branching density, curvature, and compartment shape, the errors introduced by these overlaps may or may not require attention. For many whole-neuron morphologies with thin neurite processes much longer than their radius, like the one shown in Figure 9, the contributions of segment overlaps to volume and surface area relative to the total may be negligibly small. However, if the mesh has wide compartments, high curvature, and high branching density, the overlaps can contribute substantially. In toric spines, the volume and lateral surface area estimated from segments of the cable model are consistently greater than those of the mesh because of overlaps. Summed segment volume and lateral surface area therefore include superfluous contributions where segments overlap at branch points. We estimate those contributions and subtract them when comparing the cable model to the mesh. Subtracting overlaps removes those contributions, making the geometric errors in Table 1 more interpretable.

To estimate overlaps, we use a cylinder-intersection model based on the Steinmetz solid: the shape of the intersection of two perpendicular cylinders of equal radius. Overlap for degree-2 vertices is assumed negligible: overlap on the inside of a bend is approximately balanced by missing geometry on the outside. For degree *d >* 2 vertices, each additional branch beyond degree 2 is approximated as an equal-radius branch intersecting a straight cable at right angles. For volume, that reference geometry is half of an equal-radius Steinmetz solid. The intersection volume of two perpendicular cylinders of radius *r* is ^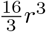^, so half of that intersection, per extra branch, contributes

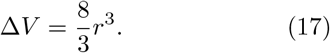

Similarly, the buried lateral surface associated with the additional branch is approximately

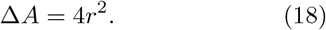

Thus, for a vertex of degree *d >* 2 and local radius *r*, the overlap corrections are

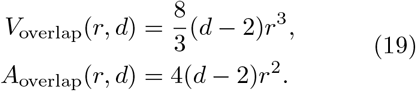

These are useful heuristics for estimating the true overlaps, which depend on various angles of intersection and non-cylindrical volumes. The scale factors 8*/*3 (volume) and 4 (area) are exposed as parameters in the `MASCAF` validation functions so that they can be adjusted if needed.

### 3.2 Simulation Validation

To verify the compatibility of generated cable models with multicompartmental simulation, reconstructed toric spine morphologies were imported into `Arbor` and simulated under standard alpha-synapse stimulation. The objective of these simulations was to confirm that morphologies produced by the `MASCAF` pipeline could be stably instantiated, discretized, and simulated without numerical or topological failure. Passive membrane parameters were assigned uniformly across compartments, and membrane potential probes were placed at every compartment to evaluate voltage propagation throughout the model.

The resulting simulations exhibited stable electrotonic membrane dynamics and continuous voltage propagation across all tested morphologies, including cyclic regions of the toric spines. With neck coordinates retrieved from imaging data, we attached a large sink volume to the spine model at that point to allow for realistic electrotonic coupling to a dendritic compartment (Koch 1999; Stuart et al. 2016). A snapshot of one such simulation is shown in Figure 11. Locations of 25 active postsynaptic densities were retrieved from imaging data and recreated with alpha-synapse models (Carnevale and Hines 2006). No simulator instability, disconnected compartments, or topology-related discretization failures were observed during model initialization or time integration. Voltage responses followed the qualitative behavior expected from passive cable theory, with attenuation and temporal smoothing increasing as a function of path length and branching complexity (Rall 1959, 1962).

**Fig. 11:**
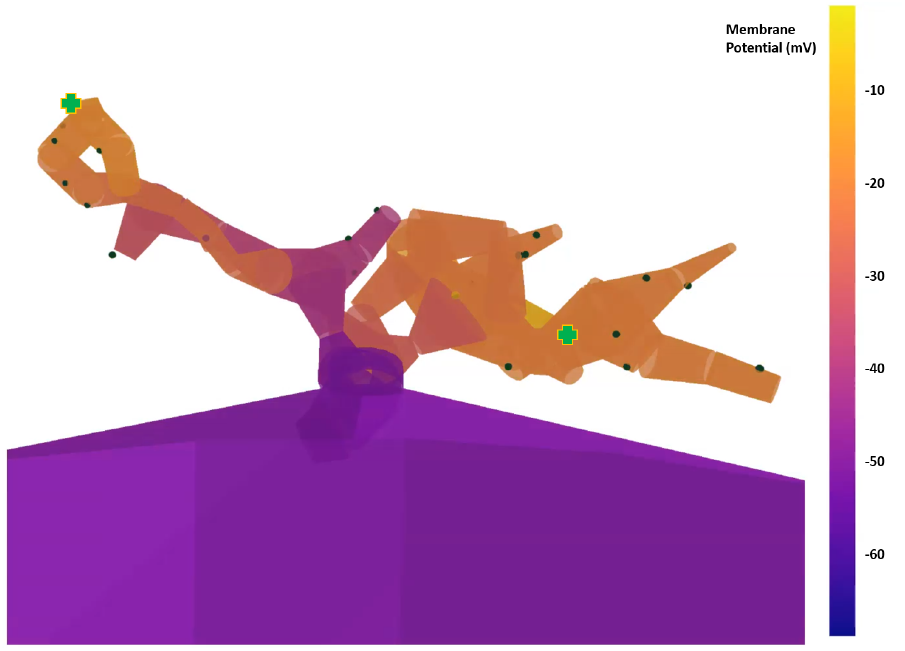
Snapshot from an `Arbor` simulation using a cable model from `MASCAF`. The spine is attached to a large sink volume to allow for realistic electrotonic loading. Membrane potential (mV) is shown on the color scale. Synapse zones (25) are rendered on the surface as dark green dots. Two locations that received a presynaptic activation are marked with green plus signs (+); their changing conductances led to the visible local depolarization.

To further validate the cable models quantitatively, we measured peak membrane potential at every compartment as a function of geodesic distance from the stimulus site, under passive membrane parameters with a single alpha-synapse stimulus. Figure 12 shows the resulting attenuation profile for toric spine #1, the morphology of which is shown in Figure 8. Peak voltage decays monotonically with path distance, consistent with electrotonic attenuation predicted by passive cable theory (Rall 1959, 1962). Furthermore, the scatter of compartment peaks at similar path distances is representative of electrotonic spread over the branching and looped cable model.

**Fig. 12:**
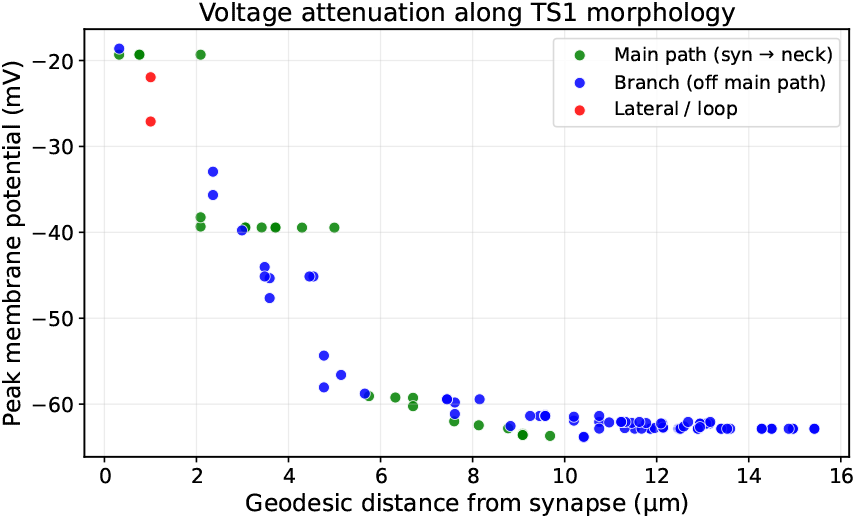
Peak membrane potential vs. geodesic distance from a distal alpha-synapse stimulus site in a passive simulation of toric spine #1 using `Arbor`. Monotonic decay is consistent with electrotonic attenuation in a passive cable model.

These results demonstrate that cable models generated using `MASCAF` are compatible with modern multicompartmental simulators and produce electrophysiological behavior consistent with passive cable theory, supporting the geometric and topological metrics reported above.

## 4 Discussion

We present the Mesh and Skeleton Cable Fitting (`MASCAF`) pipeline for fitting cable models (e.g., SWC) to 3D surface meshes, particularly suited for topologically complex neuronal structures. Cable models, like the SWC format, are the most common morphology specification used in multicompartmental simulators like `NEURON` and `Arbor`, due to their straightforward use in cable-equation models (Ascoli et al. 2007; Mehta et al. 2023). Development of `MASCAF` was motivated by the notable discovery of a new kind of dendritic spine on auditory space-specific neurons in the barn owl’s midbrain. Dubbed toric spines (Sanculi et al. 2020), they prominently feature topological holes, whose function is unknown. Cyclic topology of these morphologies breaks the assumption of tree-like neural structures (Oxford Instruments; Peng et al. 2014; Sato et al. 2000). We showed how the CGAL Triangulated Surface Mesh Skeletonization operation is used to compute midline skeletons for surface meshes via mean curvature flow. Structural cycles are retained; very small holes can be dropped in skeletonization, and the maximum edge length can remove a short cycle during resampling (Sections 2.3.1 and 3.1), so the cable-model cycle count need not equal the mesh genus. The `MASCAF` Python code (Fox 2026) uses the combined mesh and skeleton data to construct a cable model and write it to an SWC file, and also includes tools for quality optimization of the skeleton and of the morphology graph prior to radius fitting. For cyclic topologies, the output file is a valid arborescence, a rooted, directed tree, with cycle-closure directives carried in the file header; these can be read during the simulation phase to establish electrical connections, thereby preserving the real morphology in multicompartmental simulations.

### 4.1 Parameters, Limitations, & Opportunities

The most important global parameters of the `MASCAF` pipeline for cable model quality are *w*_*H*_, the `quality speed tradeoff` in `CGAL-TSMS`, and *l*, the maximum edge length in the cable fitting algorithm. As discussed in Subsections 2.2.2 and 2.2.4, *w*_*H*_ controls both the small-scale quality of the skeleton and its branching density. For small values of *w*_*H*_, the skeleton may not follow the mid-line well, while for large values, the skeleton may include spurious branching. Spurious branching describes branches in the skeleton graph *G*_Σ_ that have no corresponding volume in the mesh *M*_*S*_. This is a problem because it can introduce unrealistic structure in the cable model, but simple volumetric validation metrics might give acceptably low errors. This is a limitation of the pipeline because, in general, the user might run `CGAL-TSMS` for several values of *w*_*H*_ and choose one based on visual accuracy, since we do not have an algorithmic metric for spurious branching. However, a robust oracle system could be developed to measure geometric properties of the mesh and suggest values for `CGAL-TSMS` parameters.

The maximum edge length *l* presents a similar issue. If *l* is too small, then the number of compartments becomes large, and thus the simulation cost grows. If *l* is too large, then the model and simulation may not be a realistic representation of the biological system. A simple heuristic in the `MASCAF` Python code sets *l* to a scalar multiple of the median of the shape diameter function. It also suggests basis-optimization parameters from other geometric measurements, using settings chosen for the meshes in this study. This selection improved automation of cable model fitting for toric spines. However, a scalar function of the shape diameter is not a very robust way to choose *l*, and better parameter heuristics remain an open problem.

The basis graph optimization process described in Section 2 is under active development and, as such, the software implementation has some extended functionality that is beyond the scope of this paper. While the routine is designed to converge to a stationary configuration, it is challenging to design a robust, efficient, and convergent algorithm that centers vertices without sometimes introducing other errors. As such, the implementation described in this paper is often best used for making small global improvements to the morphology graph in a limited number of iterations (10–20).

The overlap corrections used in geometric validation (Section 3.1.1) are local heuristics based on equal-radius, right-angle cylinder intersections, half of a Steinmetz solid per extra branch, not exact Boolean unions of the frustum segments. Junctions in reconstructed morphologies are in general neither equal-radius nor orthogonal. The volume and area scale factors (defaults 8*/*3 and 4) are therefore exposed as parameters in the `MASCAF` validation functions. While this is more accurate than a simple spherical-overlap model, these default values could be tuned more carefully. An opportunity for future development of `MASCAF` is to extend the current MCFS approach with medial-axis centering methods, such as those used by TEASAR (Sato et al. 2000) and CoverageAxis (Wang et al. 2024), which are commonly used for skeletonization of 3D voxel data. While MCFS retains structural cycles, a medial-axis approach to basis graph refinement could potentially reduce errors in medial centering.

Lastly, it is important to recognize the inherent limitations of cable models as geometric representations of complex 3D shapes. Surface concavity, for example in toric spine #67, reliably leads to stubborn geometric errors, as shown in Table 1. While an accurate skeleton calculation should in principle preserve path lengths through the mesh volume, `MASCAF`’s cable fitting algorithm does not explicitly enforce this. Development of more flexible geometric representations of morphology for biophysical simulators would be a valuable contribution to this field of research.

### 4.2 Future Work

The development of `MASCAF` removes a key technical barrier in the detailed analysis of auditory space-specific neurons in the midbrain of the barn owl. High-resolution imaging data has provided unprecedented levels of detail in the anatomy and morphology of these systems, but construction of morphologically accurate simulations from that data has proven challenging. With cable-model construction from complex surface meshes now feasible, the next step involves constructing explanatory and predictive models for biophysical function of toric spines and SSNs. With `MASCAF`, we are able to perform simulations in `Arbor` of isolated toric spines with realistic synaptic and current mechanisms, probing the dynamics of signal integration. For the full SSN model, we take a multi-scale approach, grafting cable models of individual toric spines onto cable models of whole-neuron membranes. In this way we may reconstruct large and small scales at high resolution, and then probe full-scale simulations to illuminate the function of toric spines as components of a sophisticated neural computation system.

## Declarations, Code, & Data

Research reported in this publication was supported by the National Institute of Neurological Disorders and Stroke and the National Institute on Deafness and Other Communication Disorders of the National Institutes of Health under award numbers RF1NS132812 and R01DC007690. The content is solely the responsibility of the authors and does not necessarily represent the official views of the National Institutes of Health.

The `MASCAF` Python code is free and open-source: https://github.com/jmrfox/mascaf. The nine toric spine surface meshes originally prepared by Sanculi et al. (2020) and used in this study are available at https://github.com/jmrfox/mascaf/tree/main/data/mesh/processed.

## Appendix A MASCAF Pipeline Flowchart

**Fig. A1:**
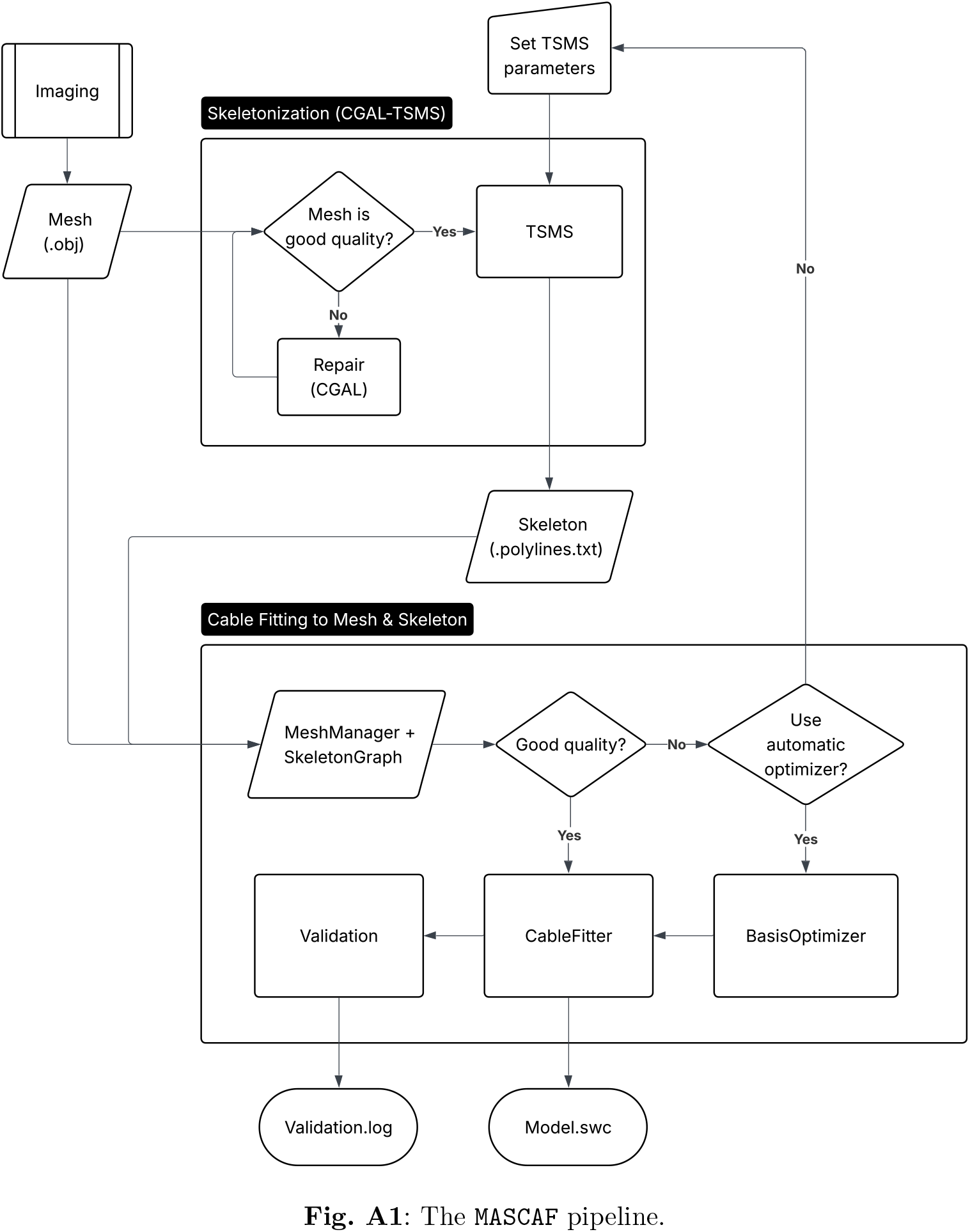
The `MASCAF` pipeline.

## Appendix B Algorithms

### B.1 Basis Optimization

#### Algorithm 1

Basis Optimization

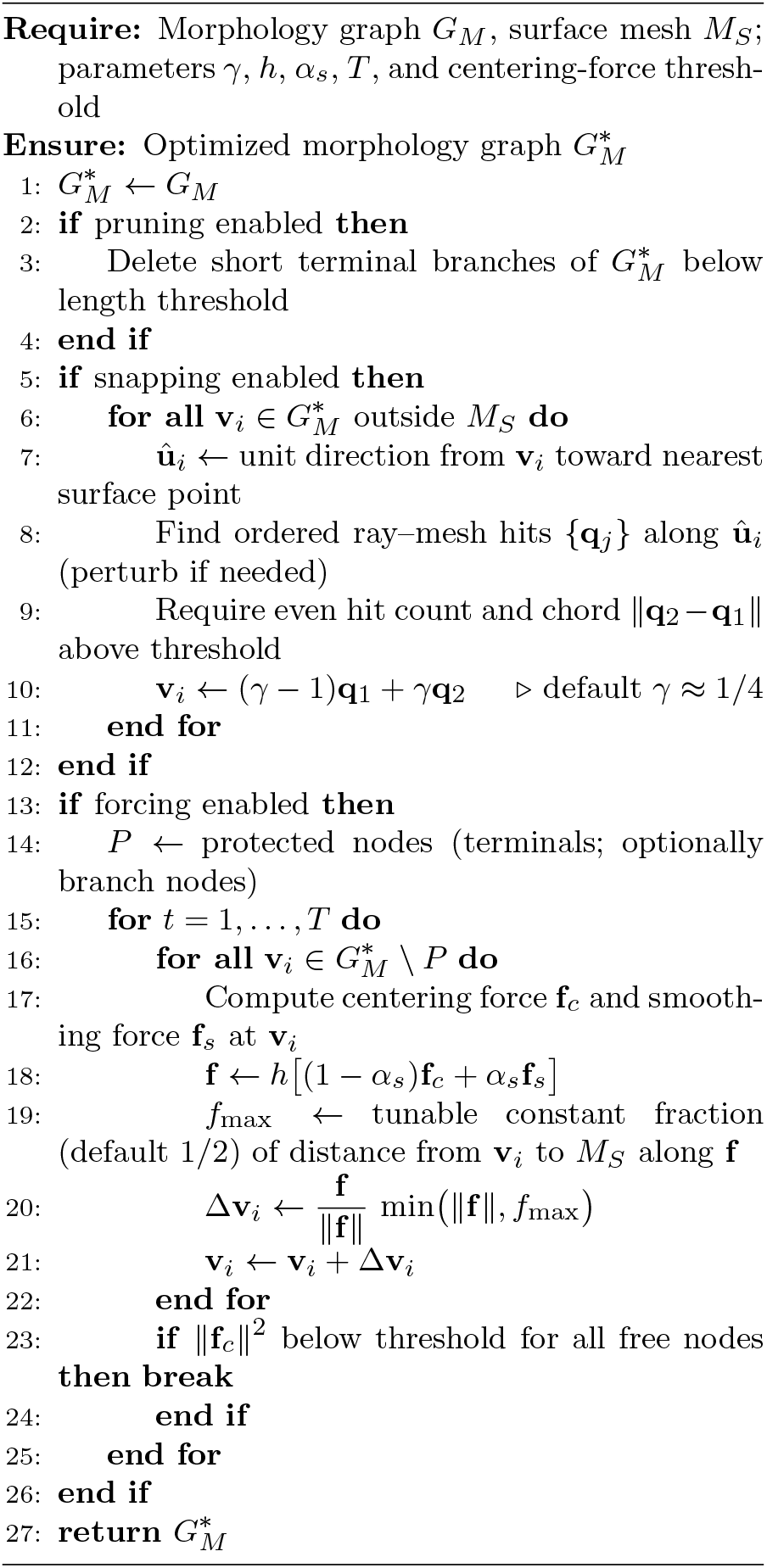

### B.2 Cable Model Fitting

#### Algorithm 2

Morphology Graph Construction and Radius Fitting

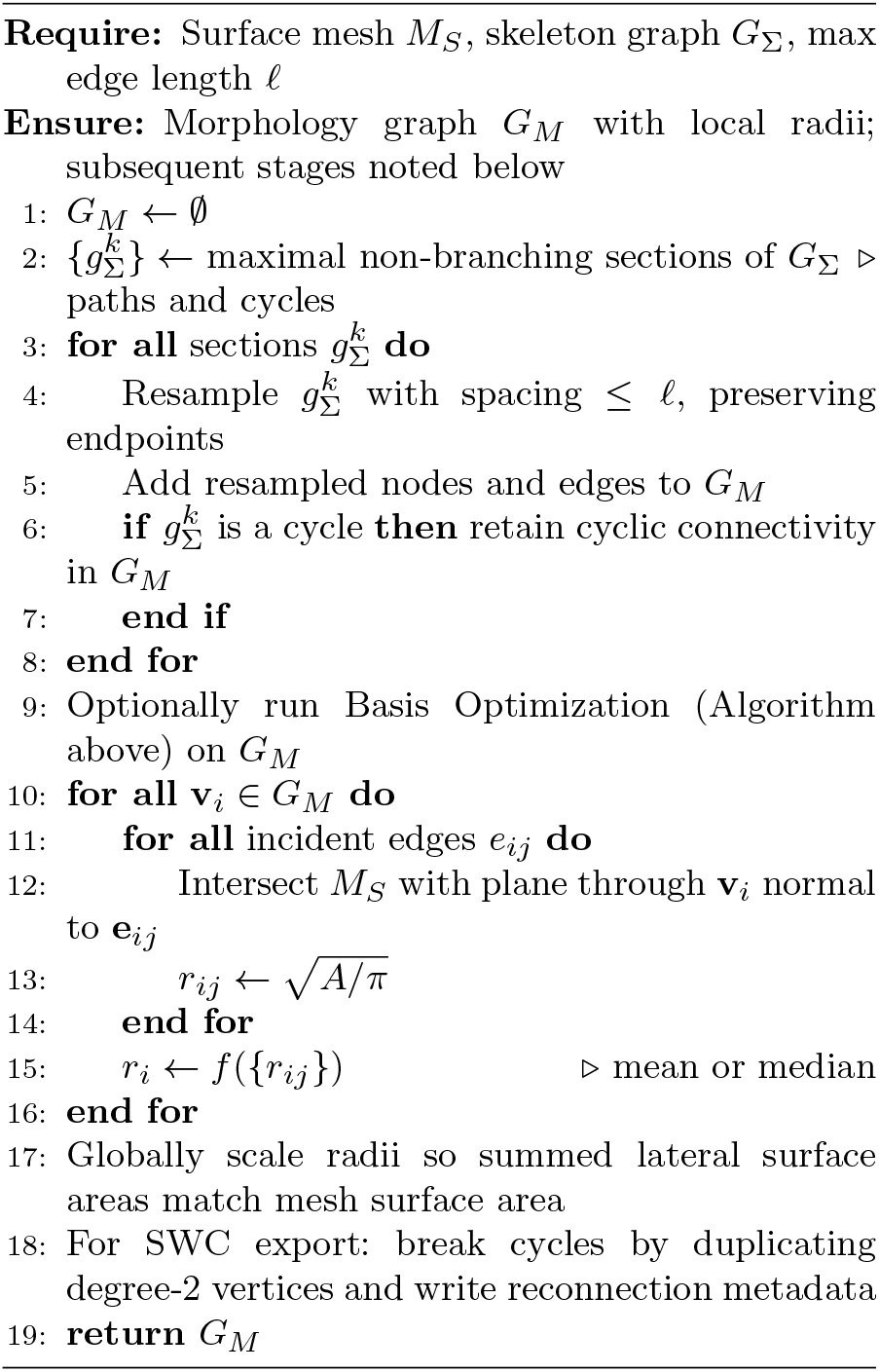

## Appendix C Related Software Tools

Fitting cable models to surface meshes is a common task in neuroscience, and there are a number of software tools that can be used to do this.

Commercial platforms such as `Imaris` (Oxford Instruments) provide powerful workflows for fitting cable models directly to imaging data and are widely used to reconstruct neuronal morphologies and connectomes from microscopy data. These tools are primarily designed for voxel-based image data and assume tree-like structures.

`Vaa3D` (Peng et al. 2014) is a multi-feature open-source program for imaging data manipulation that supports neuron reconstruction from volumetric datasets and provides cable-model outputs compatible with standard simulators. It is optimized for tracing neuronal arbors from image stacks and large-scale connectomics datasets.

Voxel-based medial-axis methods, most notably the Tree-structure Extraction Algorithm for Accurate and Robust Skeletons (`TEASAR`) (Sato et al. 2000), are related to the skeletonization approach presented here. They operate on labeled volumetric segmentations and iteratively extract centerlines by computing shortest paths to the surface in a grid. Modern implementations such as `Kimimaro` (Silversmith et al. 2021) extend this approach to large-scale dense connectomics datasets by incorporating efficient graph traversal and anisotropic distance transforms. `Kimimaro` is particularly designed to skeletonize large volumetric segmentations with high throughput and robustness. These methods are closely integrated with Google’s `Neuroglancer` (Maitin-Shepard et al. 2021) ecosystem, including tools such as Igneous (Silversmith et al. 2022) for distributed data processing and `CloudVolume` (Tartavull and Silversmith 2019) for scalable data access and visualization. These methods operate on discrete voxel grids and produce skeletons that are inherently dependent on the resolution and discretization of the volumetric segmentation. While `TEASAR`-based methods can estimate local radius using distance-to-boundary fields, they do not directly fit cable models suitable for multicompartmental simulation.

The Coverage Axis approach (Wang et al. 2024), also medial-axis-based, provides an alternative strategy for extracting skeletal representations directly from surface meshes and point clouds. These methods aim to compute a centered and well-covered medial structure by combining geometric and sampling-based criteria, and have been shown to produce high-quality skeletons for complex shapes.

`Skeletor` (Schlegel 2021) is a popular tool for mesh skeletonization and can produce quality skeletons for typical tree-like geometries, but it failed for toric spines.

Importantly, although we focus on the `CGAL-TSMS` algorithm in this paper, `MASCAF` can accept any polyline-based skeleton representation as input, regardless of the method used to compute it. This modular design allows alternative skeletonization methods, including the medial-axis approaches listed above, to be incorporated into the `MASCAF` pipeline if desired, while retaining the advantages of robust radius estimation and simulator compatibility.

## Notes

### Competing Interest Statement

The authors have declared no competing interest.

### Summary of Updates

This substantial revision, prompted by peer review, addresses many issues. These include, but are not limited to, an error analysis of geometric fits, the introduction of control cases, new diagrams illustrating optimization schemes, a computational cost report, and an appendix on comparable software.

